# Seeing light from a different angle: the effects of diffuse light on the function, structure, and growth of tomato plants

**DOI:** 10.1101/2022.02.15.480600

**Authors:** Kendra B. L. Ellertson, Gregory R. Goldsmith, Z. Carter Berry

## Abstract

While considerable attention has been paid to how plants respond to changes in the spectral distribution and quantity of light, less attention has been paid to how plants respond to changes in the angular qualities of light. Evidence from both leaf- and ecosystem-scale measurements indicate that plants vary in their response to diffuse compared to direct light growing environments. Because of the significant implications for agricultural production, we quantified how changes in light quality affect the structure, function, and growth of Roma tomatoes in an open-air greenhouse experiment with direct and diffuse light treatments. Diffuse light conditions (ca. 50-60% diffuse) were created with a glass coating to diffuse light without significantly reducing the quantity of light. We measured leaf physiology and structure, as well as whole plant physiology, morphology, and growth. Light-saturated photosynthetic rates were set by the growing light environment and were unchanged by short-term exposure to the opposite light environment. Thus, after two months, plants in the diffuse light treatment demonstrated lower photosynthesis and had thinner leaves with higher chlorophyll concentration. However, relative growth rates did not differ between treatments and plants grown in diffuse light had significantly higher biomass at the conclusion of the experiment. While there was no difference in leaf or whole-plant water-use efficiency, plants in the diffuse light treatment demonstrated significantly lower leaf temperatures, highlighting the potential for diffuse light coatings and/or materials to reduce greenhouse energy use. Our results highlight the need to advance our understanding of the effects of diffuse light conditions on agricultural crops growing on a changing planet.

## 1. Introduction

For plants, not all light is equal. The quantity and quality of light reaching Earth’s surface can have wide-ranging effects on leaf, plant, and ecosystem function (Berry & Goldsmith. 2020, Brodersen et al. 2008, Li and Yang 2015, Durand et al. 2021). It has long been recognized that the amount (quantity) and wavelengths (spectral quality) are key drivers of photosynthetic rates and plant productivity (Dueck et al. 2012, Mercado et al. 2009). However, the effects of the diffuseness of light (angular quality) on rates of photosynthesis have received less attention.

The angular quality of light can be defined as the angle of incidence of light relative to the leaf surface. Light emanates from the sun as direct, parallel beams and then becomes scattered when atmospheric particles change the direction of incoming solar radiation. As a result of scattering, some proportion of light always arrives to the canopy at a wide array of angles (Brodersen et al. 2008, Dueck et al. 2012, Mercado et al. 2009, Roderick et al. 2001, Urban et al. 2012). While the diffuse component of light can vary across locations and sky conditions, it generally ranges from 15 to 40% under clear midday conditions (Berry & Goldsmith 2020, Spitters et al. 1986, Steven 1977). For plants, light can also be scattered by the plant canopy itself or, for cultivated plants, by various greenhouse materials.

In direct light conditions, leaves at the top of a plant canopy are subjected to high light intensity while leaves in lower parts of the canopy receive less light or are completely shaded (Brodersen et al. 2008, Mercado et al. 2009, Roderick et al. 2001). In contrast, when light is diffuse, different layers of the canopy may receive light more consistently. Previous research has suggested that diffuse light can increase rates of photosynthesis, especially since light is more evenly distributed across the canopy and leaf surface (Berry & Goldsmith 2020, Brodersen et al. 2008, Dueck et al. 2012, Mercado et al. 2009, Urban et al. 2012). However, not all species respond equally to diffuse light, as studies have found both increased and decreased photosynthetic rates in response to diffuse light (Brodersen et al. 2008, Urban et al. 2012, Earles et al. 2017, Berry & Goldsmith 2020). The potential mechanisms for these responses at the leaf level, including light penetration into the leaf surface altered by anatomical changes or biochemical components that optimize carbon fixation, also remain unresolved (Earles et al. 2017, Hogewoning et al. 2012, Oguchi et al., 2011).

In addition to changes to leaf structure and photosynthetic rates in diffuse light conditions, there may also be significant effects on water-use efficiency (WUE; carbon gain through photosynthesis per unit water loss through transpiration) (Berry & Goldsmith 2020). It is possible that WUE could increase under diffuse light by having higher rates of photosynthesis while also lowering rates of water loss, as mediated by reduced leaf temperature. This may be particularly relevant for agricultural settings where minimizing water use and maximizing carbon gain is paramount to producing food in a hotter and drier world. The evidence for the effects of diffuse light on WUE are even more limited but suggest that WUE can increase in diffuse light conditions at large scales (Rocha et al. 2004). Understanding the effects of diffuse compared to direct light on plant function has implications for both basic and applied research now and given future climate scenarios.

Our objective was to compare the effects of direct and diffuse light on plant structure, function, and growth. To do so, we grew tomatoes (*Solanum lycopersicum* L.) of the cultivar “Roma” because of their global importance as a worldwide commercial crop that is commonly grown in greenhouse settings (FAO 2019, USDA 2017). Tomatoes also have a short life cycle and require significant amounts of water, which provides us with the opportunity to optimize the light environment to induce changes in structure, function, and growth (Murshed et al. 2013, Wang et al. 2015, Yang et al. 2017). We expected that diffuse light would increase photosynthesis and decrease plant water use, thus leading to higher overall WUE and growth rates, as compared to plants grown in direct light conditions.

## 2. Methods

### 2.1. Experimental Setup

To determine the effects of light environment on plant structure, function, and growth, we established a control and a treatment greenhouse in east-west orientation in Orange, California in summer 2020. We constructed 2 greenhouses measuring 7.5 m x 0.6 m with open sides and a glass roof. The glass was originally positioned at 0.38 m above plant height and was raised as the height of the plants grew over the course of the experiment. For the diffuse light treatment, we treated the glass with a diffusing paint (Redufuse, Mardenkro; Baarle-Nassau, Netherlands) that was diluted in water at a 1:6 ratio and sprayed on both sides of the glass using a paint sprayer. Paint was applied until panels measured ca. 50-60% diffuse, as described in the methodology below. The manufacturer reports no effects of treatment on the spectral quality of light, which we confirmed by quantifying the spectral distribution under the direct and diffuse chambers using a fiber optic cable connected to a CCS100 compact spectrometer (Figure S1; 350-700 nm; ThorLabs, Inc., Newton, New Jersey).

We bought 80 Roma tomato seedlings from a commercial nursery and planted each seedling in a 4.2 L pot using organic potting soil on 17 July. Forty plants were grown in each greenhouse and plants were rotated on a regular basis to minimize any effects from the position in the greenhouse. Plants were established in the greenhouses when they averaged 26.9 cm in height. Plants were fertilized with 15:9:12 N:P:K (Osmocote, Outdoor & Indoor Smart-Release Plant Food Plus, Netherlands) when planted and again in the middle of the experiment. Plants were watered to field capacity every other day, or when needed, depending on weather conditions.

### 2.2. Environmental Conditions

Temperature and relative humidity were measured continuously every 15 minutes with a shielded sensor in 4 locations in each greenhouse (U123, Onset Corporation, Bourne, MA). Photosynthetically active radiation (PAR) and the amount of PAR received as direct and diffuse light were measured continuously every 15 minutes (BF5 Sunshine Sensor, DeltaT Devices, Cambridge, England). Because only two PAR sensors were available, one was left in each treatment and sensors were rotated to each chamber on a weekly basis.

Means of temperature, relative humidity, and PAR were taken from 06:00-18:00. The diffuse light treatment was ca. 0.5°C cooler on average than the direct light treatment during the day, leading to a ca. 0.4% difference in relative humidity (Figure 1A, 1B). Total (i.e., direct + diffuse) mean daytime PAR was higher in the direct (887 ± 648 μmol mol m^−2^ sec^−1^) than the diffuse (624 ± 520 μmol mol m^−2^ sec^−1^) greenhouse, likely due to a decline in total PAR in the diffuse greenhouse in late afternoon due to some structural shading (Figure 1C). Nevertheless, diffuse mean daytime PAR was almost double in the diffuse (306 ± 238 μmol mol m^−2^ sec^−1^) compared to the direct (185 ± 99 μmol mol m^−2^ sec^−1^) greenhouse (Figure 1D). Thus, the mean daytime percent of diffuse light was 25% in the direct greenhouse and 53% in the diffuse greenhouse.

**Figure 1:**
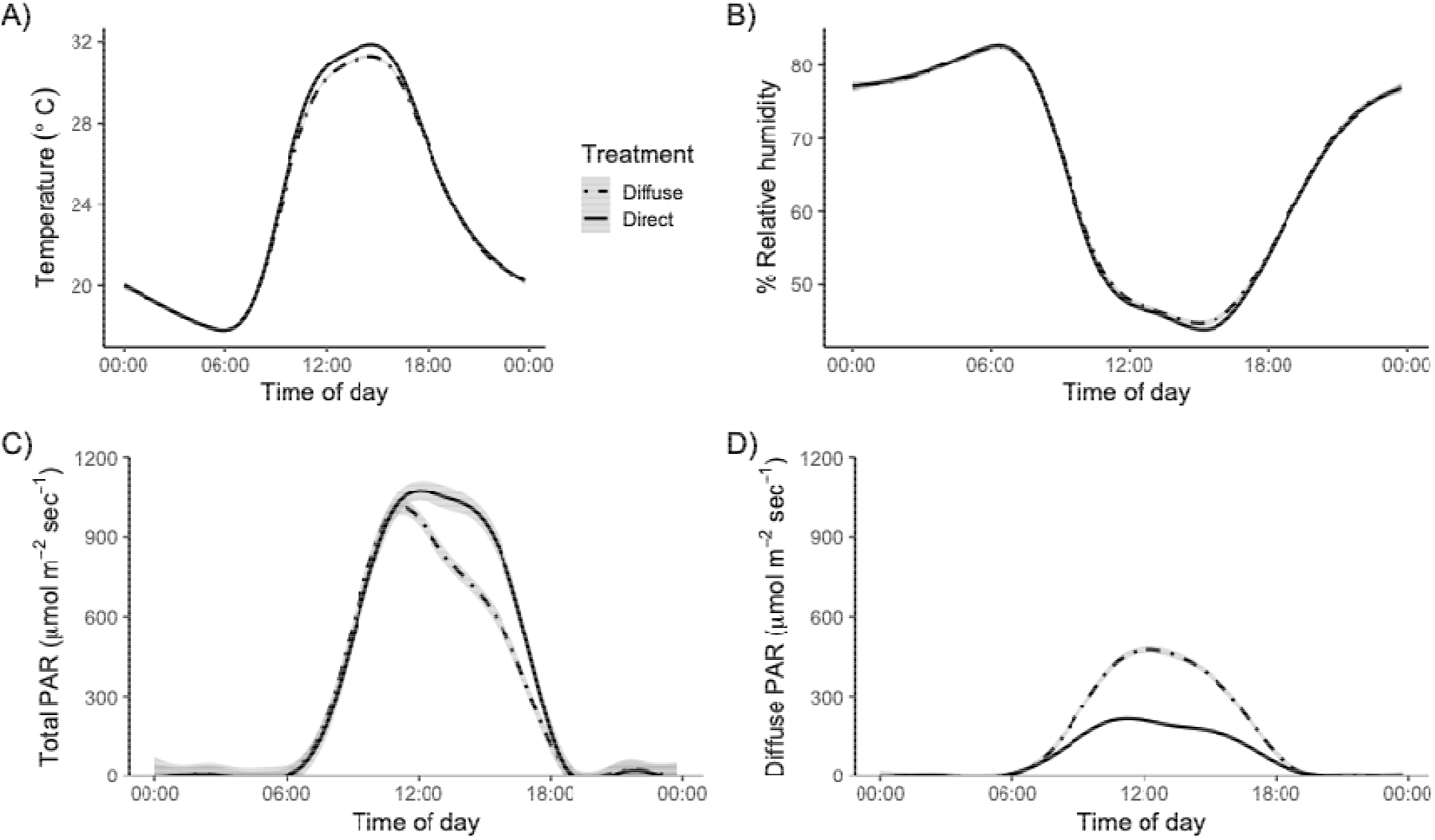
The A) temperature, B) relative humidity, C) total photosynthetically active radiation and D) diffuse photosynthetically active radiation in greenhouses with direct compared to diffuse light treatments. Data represent smoothed lines with 95% confidence intervals. Temperature and relative humidity data are from all greenhouses; photosynthetically active radiation data are from 2 sensors that were rotated weekly between greenhouses and treatments.

### 2.3. Physiological Response

We used an infrared gas analyzer (LI-6800; LI□COR Biosciences Inc., Lincoln, NE, USA) to measure photosynthesis (*A*), transpiration (*E*), stomatal conductance (*g_s_*), and intrinsic WUE (*A/g_s_*) under ambient light conditions on plants in direct and diffuse light treatments. We measured one fully mature, healthy leaf on each plant on 16 August (30 days old) and 18 September between 09:00 - 15:00 (63 days old). The leaf was placed in the 6 × 6 cm large leaf chamber (6800□13; LI□COR Biosciences Inc., Lincoln, NE, USA) and allowed to stabilize (approximately 3-5 minutes) before an instantaneous measurement was taken. The chamber air temperature was held at 28°C, relative humidity at 55%, and CO_2_ concentration at 410 ppm with a fan speed of 10,000 rpm.

### 2.4. Light Response Curves

To quantify how photosynthesis was affected by long and short-term exposure to direct and diffuse light treatments, we generated leaf photosynthetic light response curves (LRC) between 9 September and 25 September on 10 plants in each experimental treatment. To do this, we built an integrating sphere that allows us to deliver fully direct or fully diffuse light (by scattering light on ultra-white paint inside the sphere) using the existing infrared gas analyzer LED light source (Berry & Goldsmith 2020). Chamber conditions were as described above. Direct LRC were run with the LED light source directly above the leaf using photosynthetically active radiation (PAR) values at the leaf of 1290, 1113, 971, 828, 734, 663, 589, 516, 368, 183, and 16 μmol mol m^−2^ s^−1^. For diffuse LRC, the light source was moved to a 90° position on the sphere (diffuse light) and run corresponding to PAR values of 1290, 1145, 1002, 859, 751, 536, 453, 339, 226, 55, and 13 μmol mol m^−2^ s^−1^. Note that PAR values in direct and diffuse light conditions differ slightly due to the integrating sphere. At each position, the leaf was allowed to stabilize for up to 3 minutes before a measurement was taken. Light response curves were fit using Michaelis-Menten Kinetics in the DRC package for R (v3.0-1; Ritz et al. 2015).

### 2.5. Functional Traits

To analyze leaf-level response to direct and diffuse light treatments, additional functional measurements were done on 4 August. Functional traits were measured on three leaves per plant. Leaf temperature was measured with a thermocouple placed on the adaxial surface and the first stable temperature recorded. Leaf thickness was measured on each plant with a micrometer (resolution of 0.001 mm; Mitutoyo Corporation, Kawasaki, Japan). Chlorophyll content was measured using a *SPAD* handheld device (SPAD 502 Plus Chlorophyll Meter, Spectrum Technologies Inc., Aurora, IL) that was calibrated between each measurement. Leaf curling was calculated by adapting methods from Shi et al. (2007) and comparing the length and width of flattened leaves to the same measurements after leaves were allowed to curl naturally. Leaf area and specific leaf area (the ratio of leaf area to leaf dry mass) were calculated using a digital scanner and microbalance. Leaf area was analyzed using *ImageJ v. 1.51S* (National Institutes of Health, Bethesda, MD, USA).

### 2.6. Morphology

To quantify the morphological response to direct and diffuse light treatments, measurements were made approximately one month apart on 21 July, 16 August, and 18 September. Plant height was measured from the base of the main stem to the apical meristem and stem diameter was measured with electronic calipers at the base of the stem. The total number of leaves was counted manually. Relative growth rate (RGR) was calculated by dividing the difference in height or number of leaves from the start to end of the experiment by the number of elapsed days.

### 2.7. Whole Plant Physiology and Morphology

To quantify whole-plant response to direct and diffuse light treatments, whole plant biomass and WUE were measured at the end of the experiment. The night prior to measurements, all plants were watered and foil fitted around the top of the pot to prevent soil evaporation. Plants were weighed two hours before sunrise and again at sundown the same day to estimate water use. All plants were then removed from their pots, and above ground biomass was collected by clipping the stem at the soil surface. Belowground biomass was collected by gently washing soil off of roots over a 2mm sieve. Above-and below-ground biomass were dried at 60°C for at least 72 hours before being weighed. Whole-plant WUE was calculated as water uptake divided by biomass.

### 2.8. Statistical Analysis

We tested for the effects of direct compared to diffuse light treatment on different aspects of plant structure and function using t-tests. Although it may be preferable to perform an analysis with treatment, time, and their interaction where there were repeat measurements, there were insufficient observations to do so; therefore, we ran separate statistical models for each time point where appropriate. All analyses were performed in R v 4.0.3 (R Core Team, 2020).

### 2.9. Data Availability

All data will be made publicly available in the Zenodo repository upon acceptance of the manuscript.

## 3. Results

### 3.1. Leaf Physiological Response

While there were no apparent differences in leaf physiology between the direct and diffuse light treatments after one month of experimental treatment (*p* > 0.05), we did observe some notable differences after two months of treatment (Figure 2). After the second month, *A_net_* was significantly higher in the direct (14.7 ± 7.4 μmol mol m^−2^ s^−1^) compared to the diffuse (6.9 ± 2.6 μmol mol m^−2^ s^−1^) light treatment (t = −6.2, df = 48.5, *p* < 0.0001; Figure 2A). Similarly, transpiration was significantly higher in the direct (0.0067 ± 0.0047 mol m^−2^ s^−1^) compared to the diffuse (0.0039 ± 0.0022 mol m^−2^ s^−1^) light treatment after two months (t = −3.5, df = 55.0, *p* = 0.001; Figure 2B). Stomatal conductance (*g_s_*) was also significantly higher in the direct light treatment after two months (t = −3.7, df = 48.5, *p* < 0.001; Figure 2C). Given that *g_s_* increased in the direct light treatment and *A_net_* decreased in the diffuse light treatment in the second month, there was no significant difference in iWUE in the direct (47 ± 37 μmol CO_2_ mol^−1^ H_2_O) compared to the diffuse (41 ± 35 μmol CO_2_ mol^−1^ H_2_O) light treatments (t = −0.7, df = 77.7, *p* = 0.5; Figure 2D). Notably, iWUE in both direct and diffuse treatments decreased after two months.

**Figure 2:**
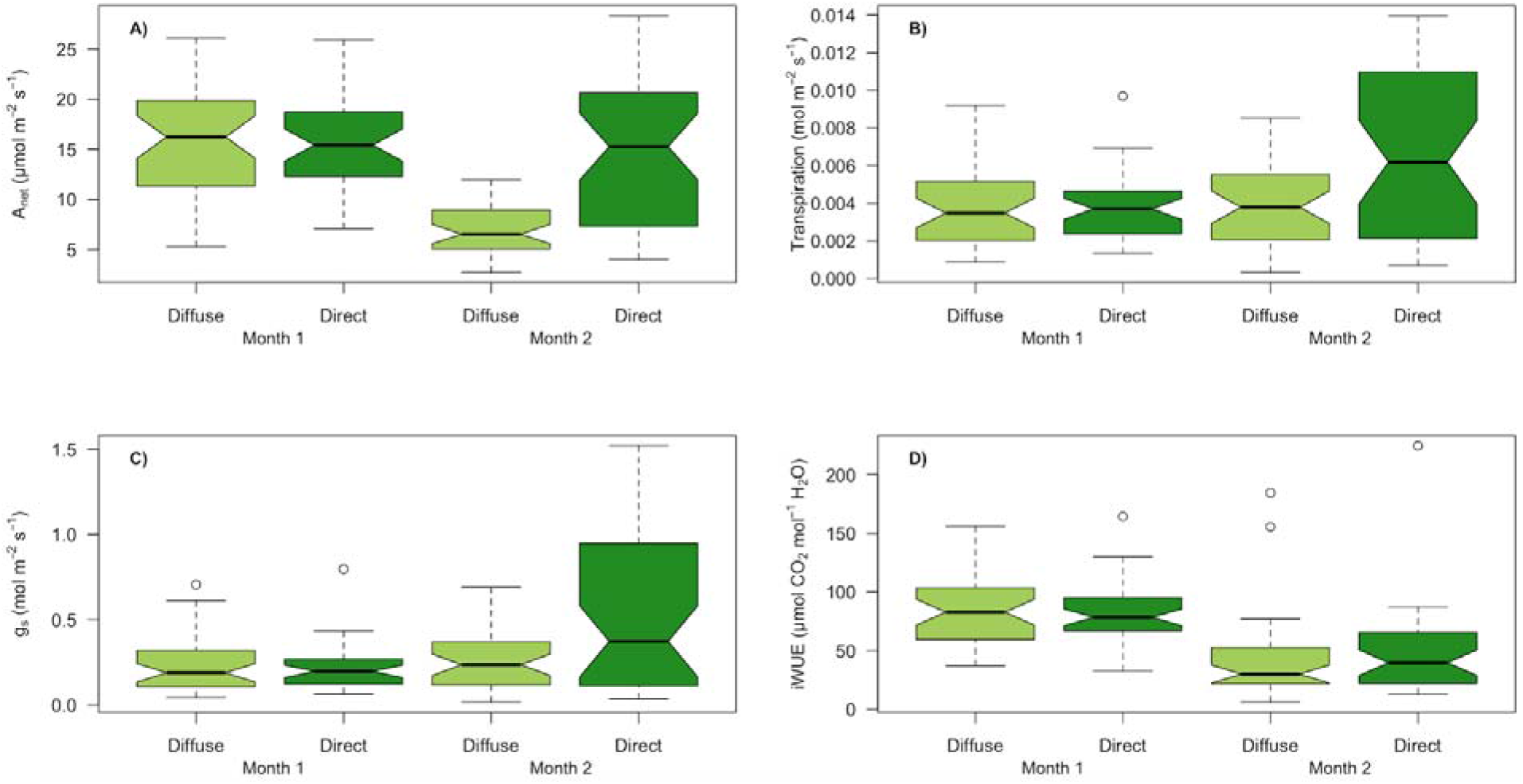
Differences in leaf physiology including A) photosynthesis (A_net_), B) transpiration, C) stomatal conductance (g_s_) and D) intrinsic water-use efficiency (iWUE) observed among tomato plants grown in direct compared to diffuse light treatments.

Light response curves of plants grown in direct light differed from those of plants grown in diffuse light. Plants grown in direct light had a greater quantum yield, maximum photosynthetic rate, and light saturation point than plants grown in diffuse light (Figure 3; Table 1). Despite being grown in distinct light environments, plants did not demonstrate distinct light response curves when the measurements were made with direct or diffuse light produced by the integrating sphere.

**Figure 3:**
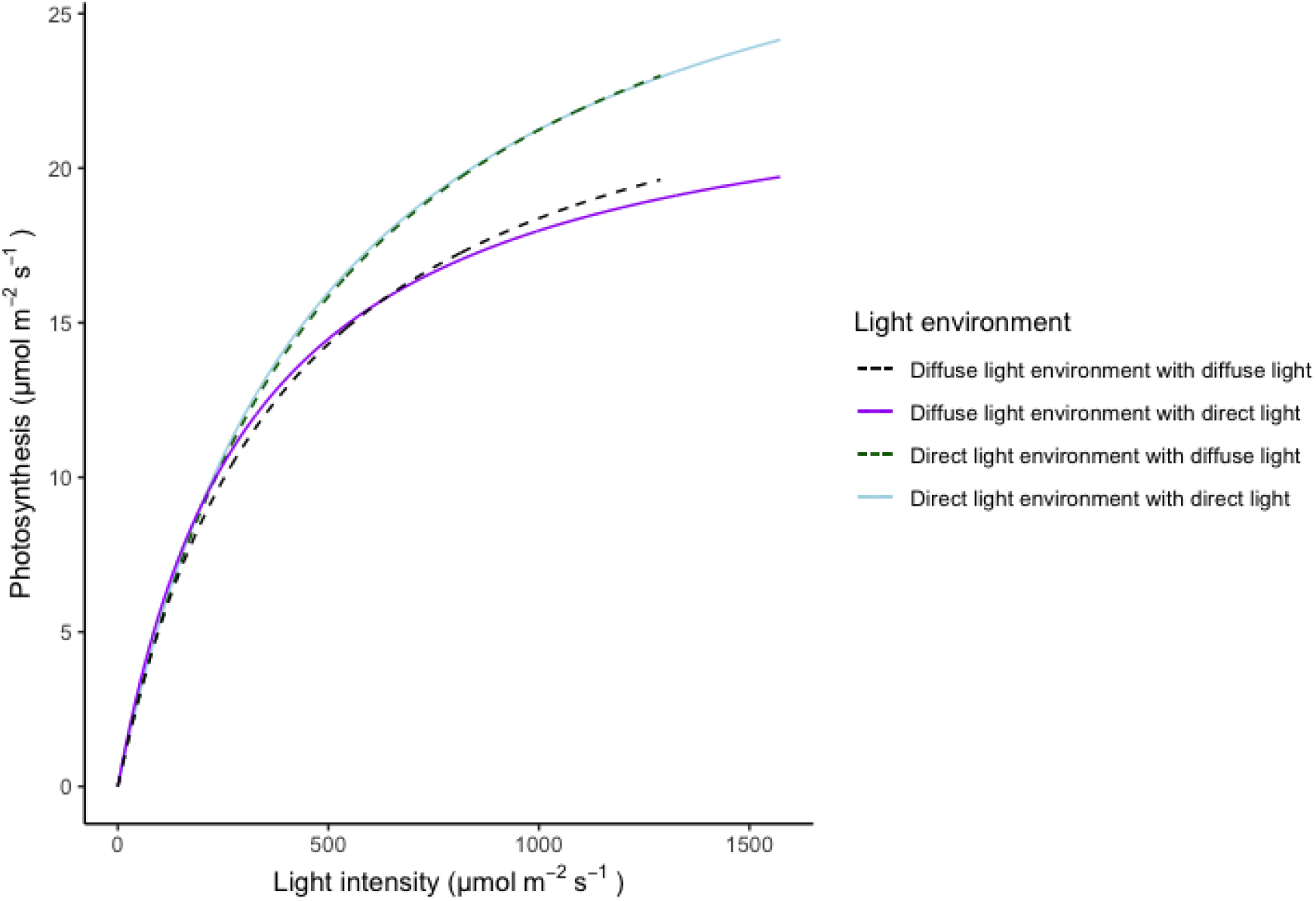
Instantaneous leaf-level light response curves observed among tomato plants grown in direct compared to diffuse light treatments. Four curves were fit from data on plants grown under either direct or diffuse light conditions and then exposed to either direct or diffuse light during the light response curves.

**Table 1:** Parameters for light response curves measured on plants in direct and diffuse light growth treatments using the integrating sphere to create direct and diffuse light. Data means ± 1 standard deviation.

| Growth treatment | Light environment | Light-saturated photosynthesis ($\mu\text{mol CO}_2 \text{ m}^{-2} \text{ s}^{-1}$) | Quantum yield ($\text{mol CO}_2 \text{ mol}^{-1}$) | Dark respiration rate ($\mu\text{mol m}^{-2} \text{ s}^{-1}$) | Light compensation point ($\mu\text{mol CO}_2 \text{ m}^{-2} \text{ s}^{-1}$) |
| --- | --- | --- | --- | --- | --- |
| Direct | Direct | $26.72 \pm 7.34$ | $0.05 \pm 0.01$ | $1.23 \pm 0.71$ | $23.18 \pm 13.71$ |
| Direct | Diffuse | $26.57 \pm 6.67$ | $0.05 \pm 0.01$ | $1.07 \pm 0.48$ | $21.23 \pm 9.85$ |
| Diffuse | Direct | $21.21 \pm 3.89$ | $0.05 \pm 0.01$ | $1.13 \pm 0.96$ | $20.06 \pm 17.927$ |
| Diffuse | Diffuse | $22.79 \pm 5.40$ | $0.05 \pm 0.01$ | $0.95 \pm 0.57$ | $19.75 \pm 13.30$ |

### 3.2. Leaf Functional Response

There were few differences in leaf functional traits between the direct and diffuse light treatments apparent after two months of experimental treatment (Figure 4). Specific leaf area differed slightly, but non-significantly, between the two treatments (*p* > 0.05; Figure 4A); however, mean leaf thickness was significantly lower in the diffuse (0.57 ± 0.12 mm) than in the direct light (0.71 ± 0.16 mm) treatment (t = 4.4, df = 74.6, *p* < 0.001; data not shown).

**Figure 4:**
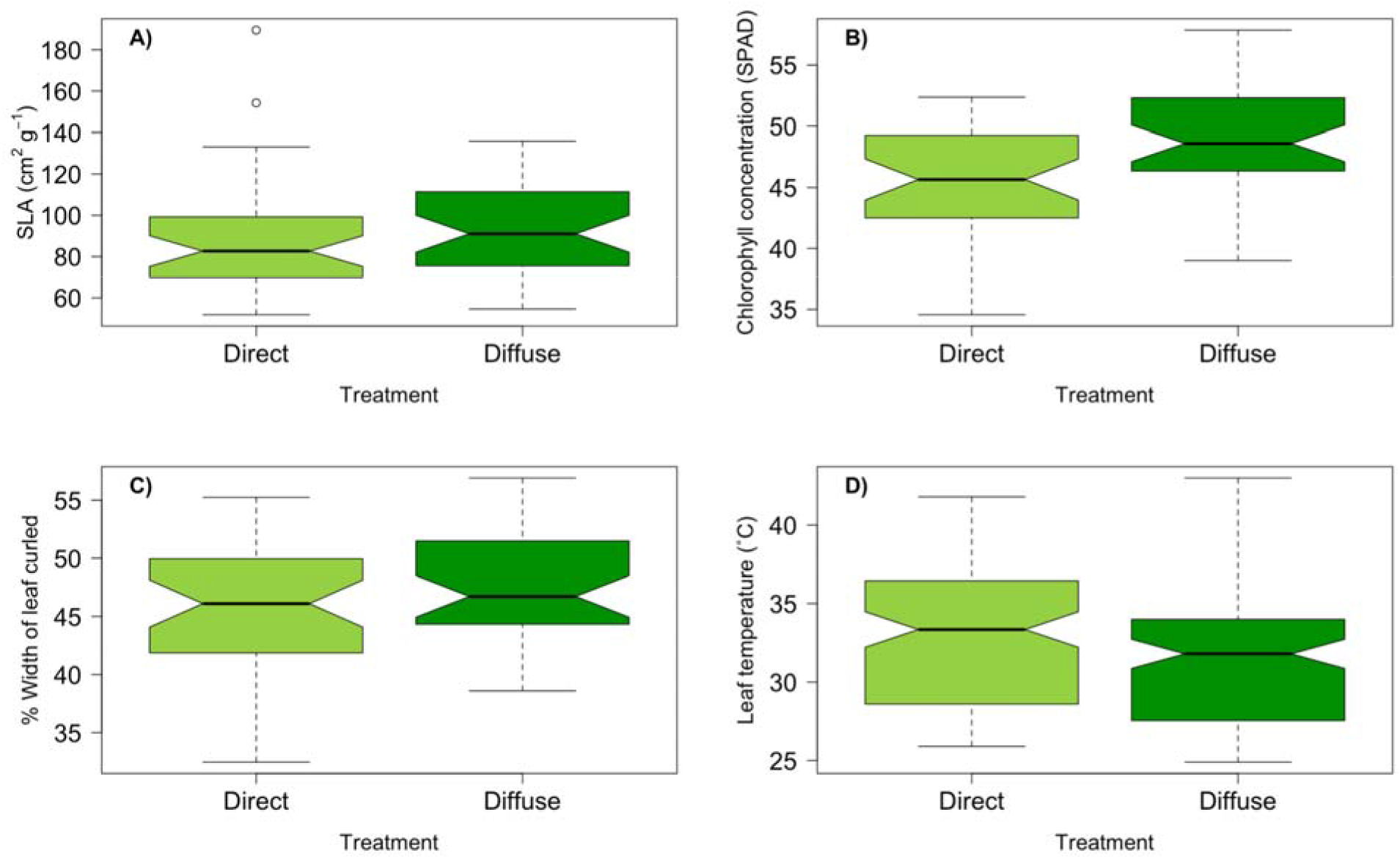
Differences in functional traits including A) specific leaf area (SLA), B) chlorophyll concentration (SPAD, C) leaf curling and D) leaf temperature observed among tomato plants grown in direct compared to diffuse light treatments after two months.

Chlorophyll content per unit leaf area was higher in the diffuse (49.1 ± 4.3 SPAD units) than the direct (45.3 ± 4.2 SPAD units) light treatment (t = −4.0, df = 78, *p* < 0.0001; Figure 4B). The average % width of leaf curling did not differ between the two treatments (*p* > 0.05; Figure 4C). We also measured leaf temperature between treatments and found that plants in the diffuse light treatment (31.2 ± 3.5°C) were approximately 2°C cooler than leaves in the direct light treatment (33.2 ± 4.4°C) (t = 3.8, df = 225.2, *p* > 0.001; Figure 4D). Overall, plants grown in the diffuse light treatment had slightly thinner leaves with higher chlorophyll content per area. Plants in the diffuse light treatment experienced lower temperatures.

### 3.3. Plant Relative Growth Rates

No differences in relative growth rates (RGR), as measured by height, stem diameter, and leaf count, were observed among plants grown in direct compared to diffuse light conditions following two months of treatment (p-value > 0.05; Figure 5A, 5B, 5C).

**Figure 5:**
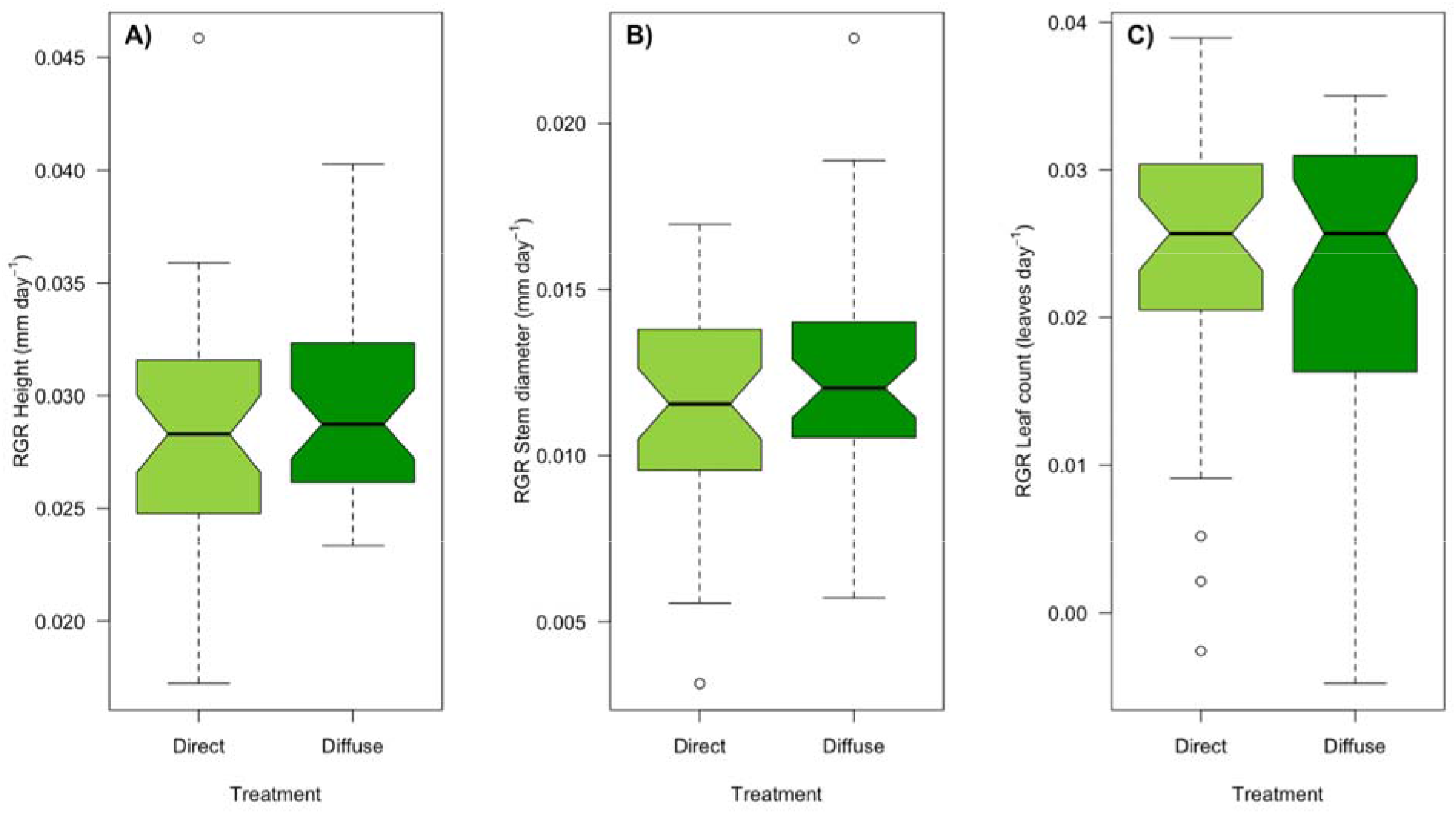
Differences in relative growth rates (RGR) for A) height, B) stem diameter, and C) leaf count observed among tomato plants grown in direct compared to diffuse light treatments after two months.

### 3.4. Plant Biomass and Whole Plant Water-Use Efficiency

Whole plant biomass and whole plant WUE were measured at the end of the experiment. By the conclusion of the experiment, the first signs of senescence were apparent in the plants grown in the direct light treatment. Plants in the diffuse light treatment had greater whole plant biomass (40.2 ± 14.1 g) than plants in the direct light treatment (28.9 ± 9.5 g) (t = −3.3913, df = 41.455, *p* > 0.01; Figure 6A) at the end of the experiment, but there was no evidence for differential allocation to above-compared to below-ground biomass between treatments. There was no difference in whole-plant WUE (p-value > 0.05; Figure 6B).

**Figure 6:**
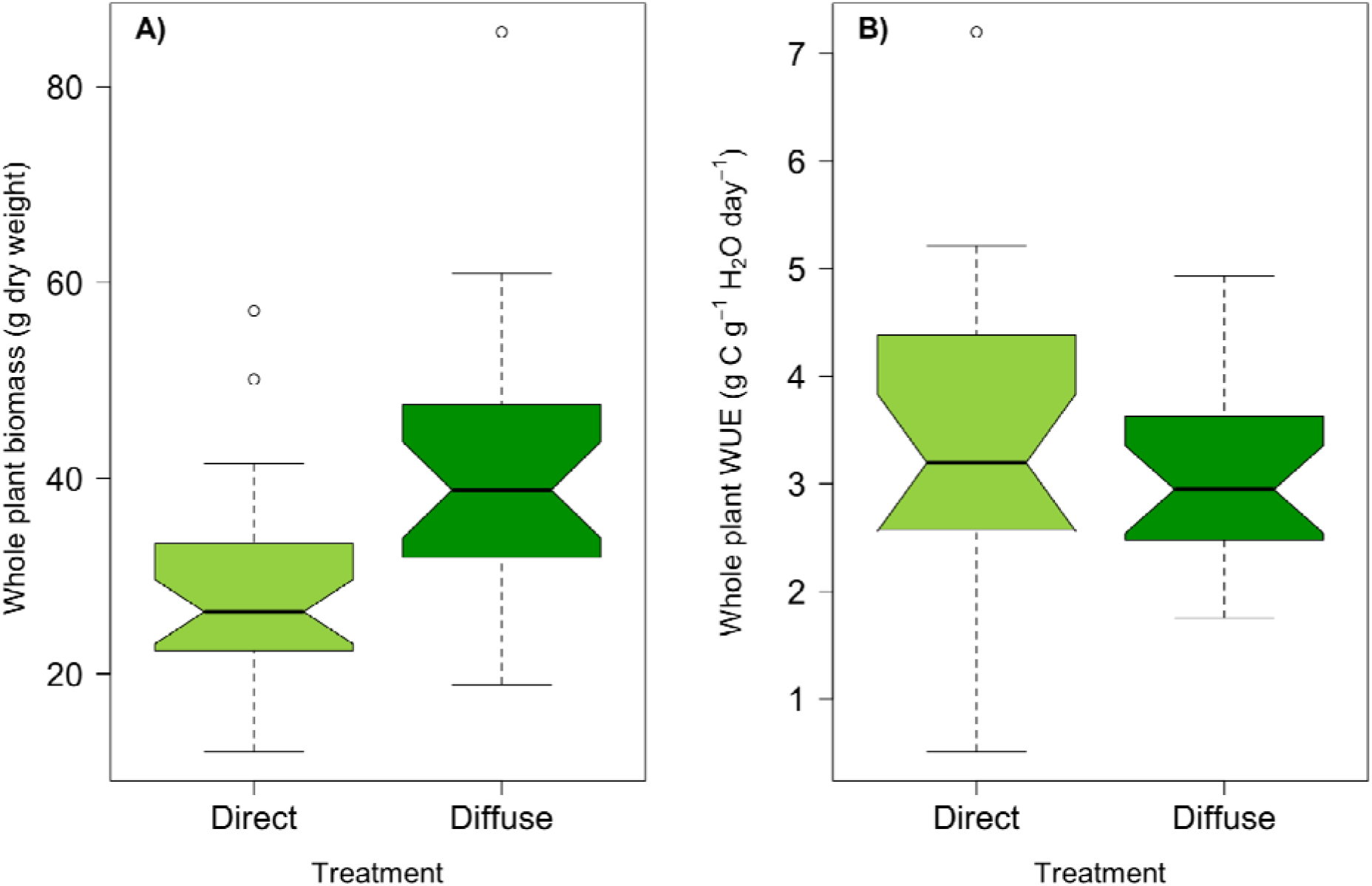
Treatment effects show that for A) Whole plant biomass (dry weight) was much greater for diffuse grown plants but for B) Whole plant WUE was greater for direct grown plants. Both measurements were taken at the end of the experiment in October after 3 months of growing.

## 4. Discussion

We compared the function, structure, growth, and productivity of tomato plants grown in direct versus diffuse growing environments. Plants in diffuse light demonstrated acclimation after two months of growth, including changes in both function (e.g., light-saturated rates of photosynthesis) and structure (e.g., thinner leaves). However, these changes did not decrease plant relative growth rates and resulted in similar (if not higher) amounts of plant biomass. Reduced photosynthesis, but higher biomass in plants grown in diffuse light may be due to differences in growth patterns (e.g., greater leaf area) or phenology (e.g., longer growth) induced by the treatment. The diffuse light environment also decreased both leaf and greenhouse temperatures, highlighting the potential for diffuse light coatings to help manage energy balance.

### 4.1. Leaf-Level Physiology

Plants in the diffuse light treatment were subject to a ca. 25% higher daytime diffuse light fraction than plants in the direct light treatment: however, we observed no differences in *A_net_*, transpiration, or *g_s_* after one month of growth (Figure 2). Only after the second month of growth did we observe a decrease in photosynthesis, transpiration, and stomatal conductance in plants in the diffuse light treatment (consistent with observations made through light-response curves). This demonstrates that photosynthetic acclimation to the diffuse light environment occurred slowly over the growth period. Ultimately, this did not lead to changes in growth rates (see discussion below) between the treatments, suggesting that photosynthetic rates and productivity were similar during the bulk of vegetative growth.

Light-saturated photosynthesis was ca. 5 μmol CO_2_ m^−2^ s^−1^ (ca. 23%) lower in plants grown in the diffuse light treatment (Figure 3). This reduction in diffuse light photosynthesis differs from Li et al. (2014) who find a 6.6% increase in whole-plant photosynthesis. They measure leaf photosynthesis in three locations and only find this increase in mid-canopy leaves. But most of their whole-plant photosynthetic increase is driven by increased light availability in the mid-canopy, not the changes in leaf physiology. Our results are more consistent with literature examining shading effects (e.g. Kläring et al., 2013), who observed a 14 – 30% reduction in diffuse light photosynthesis in tomatoes compared to direct light. Notably, short-term exposure to diffuse light in the direct light treatment, or to direct light in the diffuse light treatment, had no noticeable effect on photosynthetic light response traits (Table 1). This would indicate that growing light environment in this species governs photosynthetic traits and that those traits do not exhibit short-term plasticity in response to changes in diffuse light fraction (e.g. Berry & Goldsmith 2020).

Our results add to a growing body of research demonstrating the diverse range of responses to leaf-level physiology under diffuse light (Brodersen et al. 2008, Markvart et al. 2010, Li et al. 2014, Berry & Goldsmith 2020). Why would diffuse light lead to increased photosynthesis in some species (or even within species) and not others? The primary argument considers the physical properties of diffuse light and concluding that changes to light penetration into leaves and canopies changes the photosynthetic rate (Misson et al. 2005, Brodersen & Vogelmann 2007, Earles et al. 2017). Changes to biochemistry could also be driving these differences through differences in photosynthetic efficiency or the spatial distribution of chloroplasts within the leaf (Oguchi et al. 2011, Hogewoning et al. 2012). However, our data point to a compelling new hypothesis, that diffuse light drives changes in stomatal conductance to alter photosynthetic rates (Wang et al. 2020). The extent to which each of these hypotheses drives the photosynthetic response needs further methodical investigation.

We were equally interested in determining if diffuse light environments affected plant WUE but observed no difference in intrinsic WUE between treatments at either time point (Figure 2D). After two months of growth, plants in the diffuse light treatment demonstrated lower *A_net_*, but plants in direct light treatment demonstrated higher transpiration. These differences offset one another and there was no difference in intrinsic WUE between treatments (Figure 2D). We also observed no difference in whole-plant WUE at the conclusion of the experiment (Figure 6B), similar to the observations of tomato made by Kläring et al. (2013). As with photosynthesis, the effects of diffuse light on WUE appear to be diverse, although studies have largely focused on quantifying ecosystem-scale effects given fog or cloud cover (Baguskas et al. 2018, Knohl and Baldocchi 2008, Rocha et al. 2004). However, Knapp and Smith (1987) showed that in subalpine plants, leaf-level WUE decreased during cloud cover in some species by almost 27% or stayed relatively stable in others. A decrease in net radiation in diffuse light conditions may decrease photosynthetic rates, but also decrease water use due to changes in leaf energy balance. These results suggest that the relationship between diffuse light and plant water-carbon strategies may be context dependent. Further research on the use of diffuse light to increase WUE in agricultural applications, particularly in the context of novel greenhouse glazing materials, remains of significant interest.

### 4.2. Leaf Structure

Plants in the diffuse light treatment demonstrated significantly lower leaf thickness and higher chlorophyll content (Figure 3B). This is supported by work examining sun and shade leaves where shade leaves are typically thinner with a smaller palisade layer, but with higher chlorophyll content (Vogelmann et al. 1993). If leaf photosynthesis is driven purely by light penetration, then our diffuse light leaves should have had greater photosynthesis, but this was not the case. While we did not measure it, it is possible that there were differences in internal leaf structure by changes to proportionality of cell types (e.g., palisade vs. spongy mesophyll cells). This leaves us with an interesting result where there was a clear anatomical and morphological response to diffuse light that does not clearly link to changes in photosynthesis and transpiration. Understanding how leaf structure interacts with light penetration to drive photosynthetic rates will require further studies that simultaneously quantify variation in leaf anatomy and physiology.

Leaves on plants in the diffuse light treatment also demonstrated significantly lower temperatures than those in the direct light treatment. However, this was not clearly associated with a change in leaf energy balance as measured through transpiration rates, a change in photosynthetic rates, or a decrease in leaf curling. This is likely because tomatoes are typically grown across broad temperature ranges from 10 to 35 °C (Schwarz et al. 2014). Our data showed a leaf temperature change from 33.2°C to 31.2°C in the direct compared to the diffuse light treatment, which is well within the range of function for tomatoes. Li et al. (2014) found similar reductions in leaf temperatures and further speculated that this could minimize photodamage in diffuse light environments. For the fruits themselves, high temperatures can lead to poor fruit set, smaller fruits, and low flower numbers (Adams et al. 2001). Thus, creating growing environments with diffuse light have the potential to reduce air and leaf temperatures could lead to fruit production effects not measured here. Even a 2-3°C drop in temperature, as our results show, could decrease the energy requirements needed for large-scale greenhouse production while not compromising photosynthetic function or resultant productivity.

### 4.3. Whole-Plant Morphology

We observed no differences in plant growth rates between direct and diffuse light treatments; however, we observed higher total biomass in the diffuse light treatment at the conclusion of the experiment (Figure 6). Higher biomass in the diffuse light treatment could be a result of deeper penetration of light into the canopy (Kanniah et al. 2013, Li et al. 2014, Cheng et al. 2015) leading to greater growth even with similar or slightly lower rates of photosynthesis. This is not reflected in differences in height, stem diameter or leaf number growth rates between treatments, but could manifest as a difference in leaf area. Alternatively, we observed signs of earlier senescence among plants in the direct light treatment and believe that some biomass may have been lost. Even though Roma is a determinant variety and the date of first flowering and fruiting set did not differ between treatments (data not shown), the light environment may have altered the phenology.

In general, our results would suggest that diffuse light produces greater vegetative biomass despite no noticeable effects to standard relative growth rate measurements. This is supported by literature that find modest (2-10%) increases in diffuse light whole-plant, flower, and fruit biomass in a variety of commercially important species such as roses, chrysanthemum, anthurium, and tomato (Markvart et al. 2010, Garcia Victoria et al. 2021, Elings et al. 2012, Li et al. 2014, Holsteens et al. 2020). It should be noted that these gains in biomass have not always led to greater fruit production because of the allocation tradeoff to shoots, roots, and fruits.

## 5. Conclusion

Understanding the effects of diffuse light on plant function, structure and productivity in both field and greenhouse settings is a critical challenge for agriculture, particularly in the face of climate change (Durand et al. 2021). Diffuse light conditions will become increasing common due to changes in cloud cover and atmospheric particulate matter (Mercado et al. 2009; Roderick et al. 2001). Increased temperature and drought may also drive more agriculture into greenhouse settings, where different glazings can be employed to control the quantity and quality of radiation. Open-air, diffuse light greenhouses have the potential to reduce the energy demand for crop growth (Hemming et al. 2008; Zheng et al. 2020). We observed that diffuse light has the ability to lower leaf and greenhouse temperatures while maintaining similar light quantity, which would decrease the amount of energy spent on cooling (Elings et al., 2005).

This work, combined with the previous literature, demonstrates that there is not a unilateral response to diffuse light. In some species, photosynthesis increases while, in others, it decreases. But this does not reliably lead to predicted patterns in leaf structure or whole plant biomass. To overcome this will rely on looking past the driving hypothesis that light penetration is driving changes to diffuse light photosynthesis. An integrated framework that considers chlorophyll concentration and distribution, photosynthetic efficiency, leaf temperature effects, and stomatal responses in concert will be needed.

## Acknowledgements

We thank A. Drivas and N. Lindert for assistance in the field, B. Leahy for aiding in greenhouse construction, as well as J. Keller, L. Taylor and B. Bernardo for constructive comments. This project was funded by USDA NIFA award #2020-67014-30916 to Z.C. Berry and G.R. Goldsmith.

## Author Contributions

All authors designed the experiment. K.E. carried out the field work, analyzed the data, and wrote the manuscript with contributions from Z.C.B. and G.R.G. All authors agreed to the final version of the manuscript.

---

**Figure S1.**
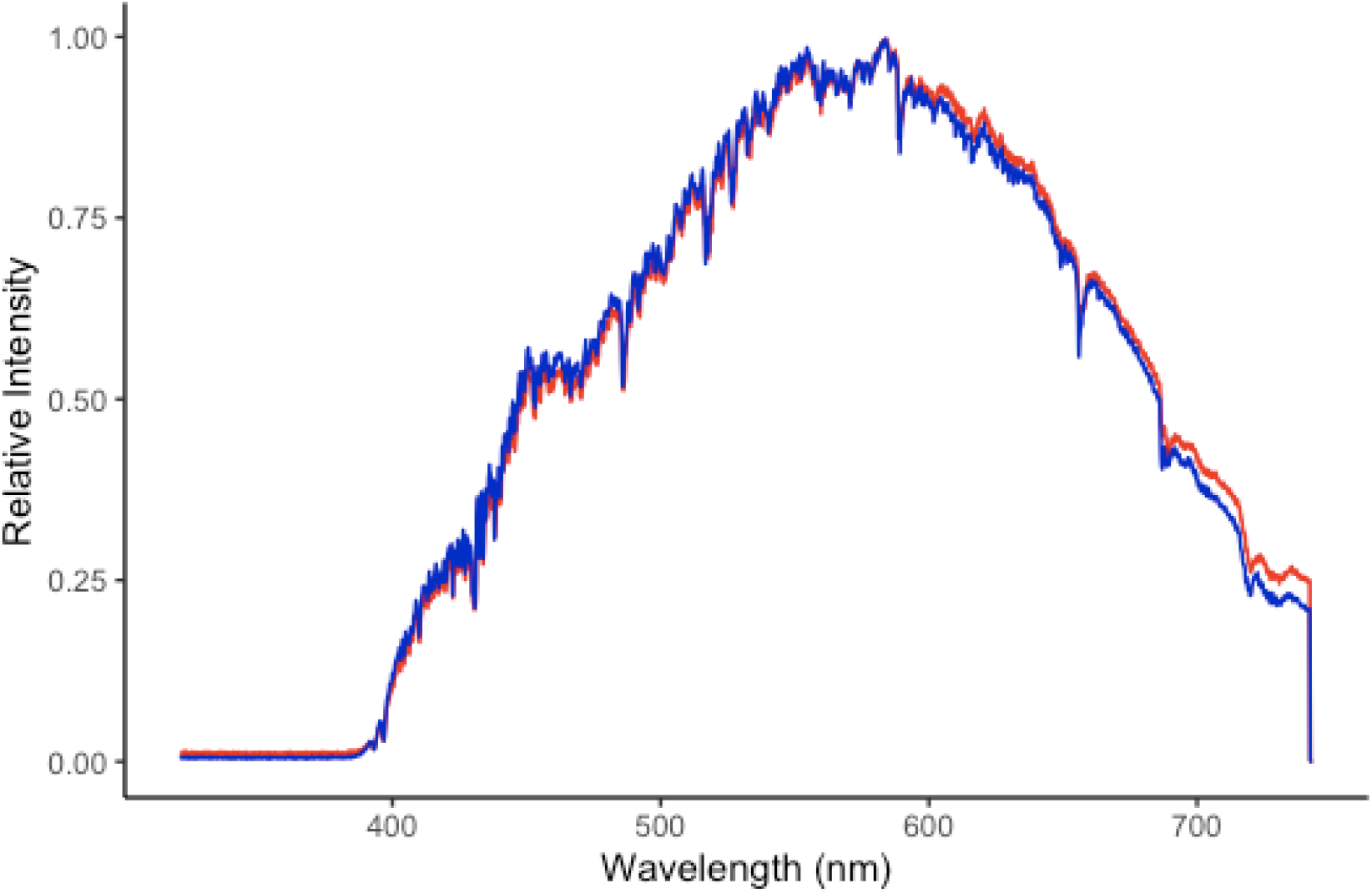
Comparison of spectral distribution of chambers with clear and diffuse paneling. The red represents clear chambers and the blue represents diffuse chambers. Measurements were made under clear sky conditions using a fiber optic cable connected to a CCS100 compact spectrometer (350-700 nm; ThorLabs, Inc., Newton, New Jersey) and data was recorded using the ThorLabs software associated with the spectrometer.

